# Do city cachers store less? The effect of urbanization and exploration on spatial memory in individual scatter hoarders

**DOI:** 10.1101/377143

**Authors:** Megan Joy Thompson, Julie Morand-Ferron

## Abstract

Urbanization has been shown to affect a variety of traits in animals, including their physiology, morphology, and behaviour, but it is less clear how cognitive traits are modified. Urban habitats contain artificially elevated food sources, such as bird feeders, that are known to affect the foraging behaviours of urban animals. As of yet however, it is not known whether urbanization and the abundance of supplemental food during the winter reduce caching behaviours and spatial memory in scatter hoarders. We aimed to examine individual variation in caching and spatial memory between and within urban and rural habitats to determine i) whether urban individuals cache less frequently and perform less accurately on a spatial task, and ii) explore, for the first time in scatter hoarders, whether slower explorers perform more accurately on a spatial task, indicating a speed-accuracy trade-off within individuals. We assessed spatial memory of wild-caught black-capped chickadees (*Poecile atricapillus*; N = 96) from 14 sites along an urban gradient. While the individuals that cached more food in captivity were all from rural environments, we find no clear evidence that caching intensity and spatial memory accuracy differ along an urban gradient, and find no significant relationship between spatial cognition and exploration of a novel environment within individuals. However, individuals that performed more accurately also tended to cache more frequently, suggesting for the first time that the specialization of spatial memory in scatter hoarders may also occur at the level of the individual in addition to the population and species levels.

## INTRODUCTION

Urbanization is occurring globally at a dramatic rate, and in response some species are declining while others are thriving in urban environments (Sih et al. 2011; Lowry et al. 2013). Species that colonize urban environments may have access to more food sources compared to rural environments, especially in variable or seasonal periods of low food availability (Lepczyk et al. 2004; Lowry et al. 2013; Tryjanowski et al. 2015). Large quantities of nutrients and energy are added into urban systems each year through commercial bird feeding activities and (Galbraith et al. 2015), as a result, urban areas are reported to support significantly more overwintering birds (Clergeau et al. 1998; Marzluff et al. 2001; Tryjanowski et al. 2015). Supplemental bird feeding has been shown to decrease over-winter mortality and thus has direct fitness consequences (Brittingham and Temple 1988; Desrochers et al. 1988). There is also evidence that evolutionary change in response to bird feeding activities can occur on short time scales as a recent study found that bill length of a common garden bird, the great tit *(Parus major)*, increased significantly in response to supplemental feeder use as a part of a 26-year study (Bosse et al. 2017). In response to predictable year-round food availability, ecological and behavioural traits of urban species are changing, especially those associated with foraging (Lowry et al. 2013; Galbraith et al. 2015).

Scatter hoarders rely on spatial memory to recall locations of many previously stored food items, which may be critical for survival through periods of food scarcity (Krebs 1990; Mcnamara et al. 1990). Scatter hoarding birds have been empirically shown to possess enhanced spatial memory and related neurological features, which they use to recall and retrieve their stored caches (reviewed in Brodin, 2010; Pravosudov & Roth, 2013). The adaptive specialization hypothesis (ASH) predicts that hoarders occupying harsher or food-scarce environments should exhibit superior spatial cognition and the corresponding neurological features to support a higher reliance on caches in their environment (Krebs 1990). ASH has been supported at both latitudinal and elevational gradients within species (Pravosudov and Roth 2013). These studies show that individuals occupying higher latitudes or elevations (harsher, food-scarce environments) cache more frequently and show superior spatial accuracy in comparison to conspecifics who occupy low latitudes or elevations. If urban areas have more abundant food sources and are less seasonally harsh environments (Lowry et al. 2013; Tryjanowski et al. 2015), urban scatter hoarders may rely less on caching behaviours and spatial memory. The selective pressures acting on specialized spatial memory may therefore become relaxed in urban scatter hoarders and the spatial abilities of these individuals may become inferior to rural conspecifics as a result. Despite artificially increased food availability via the use of feeders in developed and urban areas, spatial abilities have only recently been examined in scatter hoarders occupying urban environments (Kozlovsky, Weissgerber, & Pravosudov, 2017).

In addition to cognitive traits, certain personalities have been suggested to facilitate colonization of urban habitats (Miranda et al. 2013). Between-individual differences in cognitive abilities may cause variation in behavioural tendencies or vice versa, and therefore personality and cognition may together be affecting the responsiveness of individuals to environmental change (Griffin et al. 2015). For example, individuals with fast exploratory personalities can discover new foraging opportunities more readily (Herborn et al. 2010; van Overveld and Matthysen 2010), which may be important for colonizing urban environments (Atwell et al. 2012; Sol et al. 2013). However, it has been hypothesized that a trade-off exists between exploratory personality and quality of spatial information acquired (Verbeek et al. 1994; Sih and Del Giudice 2012). Fast explorers are assumed to collect shallow spatial information due to rapid movement through their environment (fast-but-superficial exploration) in comparison to slow explorers, who would have more gradual searching strategies and collect higher quality spatial information (slow-but-thorough exploration). There is indirect support for this hypothesis in mountain chickadees (*Poecile gambeli*) where high elevation chickadees showed superior spatial accuracy (Freas et al. 2012), and different individuals from the same site were found to also be slower explorers in comparison to low elevation chickadees (Kozlovsky et al. 2014). These findings suggest that selection for superior spatial memory in harsh environments may be directly or indirectly linked to selection for slow exploratory personality (Kozlovsky et al. 2014). However, the hypothesized trade-off between personality and cognitive accuracy has received little empirical attention in relation to spatial cognition (but see Bousquet, Petit, Arrivé, Robin, & Sueur, 2015; Schuster, Zimmermann, Hauer, & Foerster, 2017), and has never been examined at the individual-level in scatter hoarders.

This study aimed to examine ASH in scatter hoarders along an urban gradient, and explore the relationship between spatial memory and exploratory personality within-individuals. We hypothesized that stable food sources in urban areas, especially over harsh winter months, would cause urban scatter hoarders to be less reliant on food caching behaviours, and therefore decrease selective pressures on spatial memory. We predicted that urban scatter hoarders would cache less and show inferior spatial memory accuracy in comparison to their rural conspecifics. We also hypothesized a negative within-individual covariation between exploratory personality and spatial accuracy, which would be indicative of a trade-off between speed of exploration and quality of information collected.

## METHODS

### Study species, sites, and captivity

To test our hypotheses, we use the black-capped chickadee (*Poecile atricapillus*), a scatter hoarding, non-migratory passerine bird that can be found in a variety of different habitats in North America, including both urban and rural environments (Smith 1991; Foote et al. 2010). Capture of wild black-capped chickadees occurred in the years 2015 and 2016 between September 22 - December 07 at fourteen sites that varied in degree of urbanization (urban = 7, rural = 7). Urban sites were partially forested city parks surrounded by residential buildings and rural sites were completely forested areas (at least 500×500 m) > 25 km away from downtown, with captures occurring at least 300 m away from the nearest residence. All sites were at least 2 km apart to ensure that home ranges of chickadees did not overlap (10-20 ha home range; Smith 1991). Urban scores were generated for each site using remote sensing via satellite imagery to quantify the number of pixels for land cover components of forest, bare earth, tarmac, and buildings within a 1 km radius of capture locations at each site (Thompson et al. in press). These variables were used in a principal component analysis (PCA) to generate unique scores for each site for use in further analyses. Urban scores were also generated using a 200 m radius for comparison since 1 km may be over-representing the home range of chickadees (8.8-22.6 ha; Smith, 1991).

Upon capture, individuals were weighed, measured, and aged as either juvenile (hatch-year) or adult (after-hatch-year) birds by shape and wear of the outermost retrices (Pyle 1997). Birds were transported to captivity and housed in individual cages (40×60×40 cm) located in indoor aviaries that allowed auditory contact, but prohibited both physical and visual contact, and were maintained on a 10D:14N light cycle at 20 ± 1°C. Individuals could store food freely in 28 potential cache sites in their home cages. Each individual’s cage caches were counted and removed twice daily to quantify caching intensity. Each individual’s cage connected directly to a larger testing room via an opaque sliding door. We used light manipulation to control the movement of individuals between their home cages and the larger testing room to avoid unnecessary handling (Pravosudov and Clayton 2002).

Birds were housed in captivity for 5 days so individuals could be tested only a couple days after capture from the wild. While this approach is standard in the personality literature (e.g. Dingemanse et al. 2002), animals are often allowed a longer familiarization period in cognition studies, with the assumption that baseline corticosterone (stress) returns to normal levels after a number of weeks in the captive environment (Dickens, Earle, & Romero, 2009; Pravosudov, Mendoza, & Clayton, 2003). However, captivity has also been shown to affect the phenotype of wild-caught animals (Lattin et al. 2017). In particular, both neurogenesis within, and the volume of, the brain region responsible for spatial cognition in scatter hoarders (i.e. hippocampus) has been reported to decrease in response to captive conditions, and it is unknown how this may affect spatial performance (Roth et al. 2010). By testing birds a couple of days after capture, we aim to avoid potentially altering the phenotype of individuals and to increase power by testing more individuals, while controlling statistically for individual stress levels after exposure to captivity. On the fifth day in captivity, we extracted blood samples from the brachial vein of all individuals to determine sex and captive baseline corticosterone levels (stress; Pravosudov, Kitaysky, Wingfield, & Clayton, 2004). Blood samples were taken ~ 45 hrs after spatial memory tasks (~94 hrs after capture) as this was the earliest moment in which samples could be taken without disturbing individuals before behavioural assays. Baseline corticosterone levels (x = 3.88 ng/mL, range = 0.44 – 13.74) fell within reported ranges for this species under both long-term captive (Pravosudov et al. 2004) and wild (Montreuil-Spencer 2017) conditions.

Tasks presented in captivity required individuals to be able to remove pompoms (1.5 cm diameter white cotton balls) out of holes (cache sites; 1×1 cm) in search of a hidden sunflower seed reward. Therefore, a gradual behavioural shaping procedure was administered in individual cages during the morning of the second day to familiarize birds with the concept of hidden seed rewards and the motor movement required to remove pompoms. Birds were considered as trained when they removed a pompom and retrieved a seed reward from a cache site in three consecutive trials during initial training. During the afternoon of the second day, birds were further trained to search for a single food reward for a 3-hour period when not all pompom removals were rewarded.

### Exploration in a novel environment

On the third day in captivity between 7:30-11 hours, exploration behaviours of individuals were quantified using a novel environment exploration assay (Verbeek et al. 1994) within the larger testing room. Similar to Quinn et al., (2009), the duration of flights, duration of hops (number of hops*0.5 sec), number of visits to each tree (1-4), and number of visits to other features in the room (ceiling, floor, walls) were recorded live by the same observer. These variables were used in a principal component analysis (PCA) to generate a score that represented an individual’s exploration behaviour during the assay. Exploration scores were found to be significantly repeatable in a second altered environment assay (R = 0.47, CI = 0.41 – 0.51, χ^2^= 17.3, *P* < 0.001; Thompson et al. in press), and fell within the reported range for exploration behaviours (Bell et al. 2009).

### One-trial associative spatial memory task

A one-trial associative spatial memory task (Clayton and Krebs 1994) was conducted on the third day in captivity between 12:00-13:00. This task took place in the flight room using the four trees used for novel environment exploration, which each bird had been familiarized with for 30 min prior. This task included two phases. During the preliminary phase, birds entered the testing room and were given a maximum of five minutes to find and contact a specific rewarded site selected by the experimenter among 60 possible cache sites. The rewarded site contained visible sunflower seeds and was kept the same for all birds to allow for individual-level comparisons. Once a bird contacted the sunflower seed in the first phase, it was allowed 10 seconds to feed on the seed before the lights were turned off and it was returned to its home cage. Individuals then underwent a 30 min retention interval before re-entering the testing room for the experimental phase. At this time, the trees in the testing room were switched to ensure birds were not relying on visual cues from the trees and all 60 cache site contents were concealed using pompoms. During the experimental phase, birds re-entered the testing room and attempted to find the location where they had found food in the previous phase (Figure S1). Birds were food deprived for 30 min prior to the preliminary phase and throughout the task for a total of 1 hr. How quickly the birds completed the task was measured as the latency to contact the hidden seed during the experimental phase. Additionally, an individual’s spatial memory accuracy was inferred from the number of pompoms pulled (errors) before contacting the hidden seed. Individuals that did not contact the seed reward within the testing duration were excluded. Baseline corticosterone levels were not found to significantly differ between participating and non-participating individuals by habitat type (Table S1). In 2016, we assessed an individual’s spatial performance again in a second session in an attempt to assess repeatability of spatial performance. The second session occurred the next day at the same time and followed the same procedure as the first session, however the seed reward was placed in a different location.

### Statistical analysis

We evaluated caching intensity by fitting the total number of caches made in captivity with a generalized linear mixed-effects model (GLMM) under a negative binomial distribution to account for overdispersion and aggregation in the count data (O’Hara and Kotze 2010; Harrison 2014). We tested an interaction between urban score and the number of errors made during the spatial task, controlled for baseline corticosterone, and included sites as random intercepts. To include more individuals and further examine the effect of urbanization, we also fitted a model excluding errors in the spatial task as we did not have data for all individuals in this test.

Latency to contact the hidden seed (speed) in the one-trial spatial task was log-transformed to improve normality and fitted using a linear mixed-effects model (LMM). The number of errors (accuracy) within the spatial task was fitted using a GLMM with a negative binomial distribution accounting for overdispersion. In both models, we tested an interaction between urbanization and exploration score, controlled for testing order and baseline corticosterone, and included capture site as a random intercept. No difference in spatial memory performance was found between years so we pooled data over both years. Baseline corticosterone level after exposure to captivity was found to be non-significant in all spatial task models (all *P* ≥ 0.8; Table 2b) and, as we did not have data for all individuals, our final models excluded this variable in order to include the full dataset.

In an additional analysis, we also included the second session of the one-trial spatial task for individuals tested in 2016 to assess repeatability of spatial memory performance. We evaluated the number of errors (GLMM negative binomial) and latency to contact the reward (LMM) as done previously. An interaction between urbanization and exploration scores, as well as session (1 or 2) and testing order were included as fixed effects in the models. Site levels were returning zero variance when included as a random effect for both models, we therefore only controlled for individuals as random intercepts to account for pseudoreplication. We evaluated individual consistency of spatial memory accuracy and latency to contact the reward by calculating adjusted repeatability (Nakagawa and Schielzeth 2010; Griffin et al. 2015).

Continuous fixed effects in mixed models were standardized via grand mean-centering prior to analysis to improve convergence (Pinheiro and Bates 2000; Bolker et al. 2009). All tested interactions were found to be non-significant and were therefore removed so that the main effect of predictors could be evaluated. Since our data were skewed and showed overdispersion, we evaluated the robustness of each of our analyses by removing outliers that were more than two standard deviations from the mean (Miller 1991). GLMMs were fitted with a Laplace maximum likelihood approximation. We evaluated significance of fixed-effects using Type II ANOVAs and interacting terms using Type III ANOVA. We evaluated terms in Gaussian models using F-tests and non-Gaussian models using Wald chi-square tests (Bolker et al. 2009). All statistical analyses were conducted using R v.3.4.3 (R Core Team 2018). Our mixed-effects models were generated using the *lmer* and *glmer.nb* commands in the lme4 package (Bates et al. 2015).

## RESULTS

### Caching intensity

The number of cage caches was not significantly predicted by an interaction between urbanization and the number of errors during the spatial task (Table 1a) and therefore this effect was dropped. The number of errors an individual made on the spatial memory task was significantly associated with the number of cage caches (Table 1b) showing that more spatially accurate individuals cached more frequently while in captivity (Figure 1a). However, this association was non-significant when excluding observations greater than two standard deviations from the mean (all observations > 22 caches and > 33 errors, GLMM: *N* = 71, errors estimate ±SE = -1.00 ±0.95, χ^2^ = 1.10, *P* = 0.29). Urbanization did not significantly affect caching intensity when using the spatial task dataset (Table 1b), but was found to be significant when also including individuals that did not participate on the spatial task, with urban birds caching fewer food items than rural birds (Table 1c; Figure 1b). However, this effect was found to be driven mainly by extreme values and was non-significant when excluding all observations greater than two standard deviations from the mean (observations > 22 caches, GLMM: *N* = 130, urban estimate ±SE = -0.19 ±0.35, χ^2^ = 0.32, *P* = 0.57).

**Fig.1.**
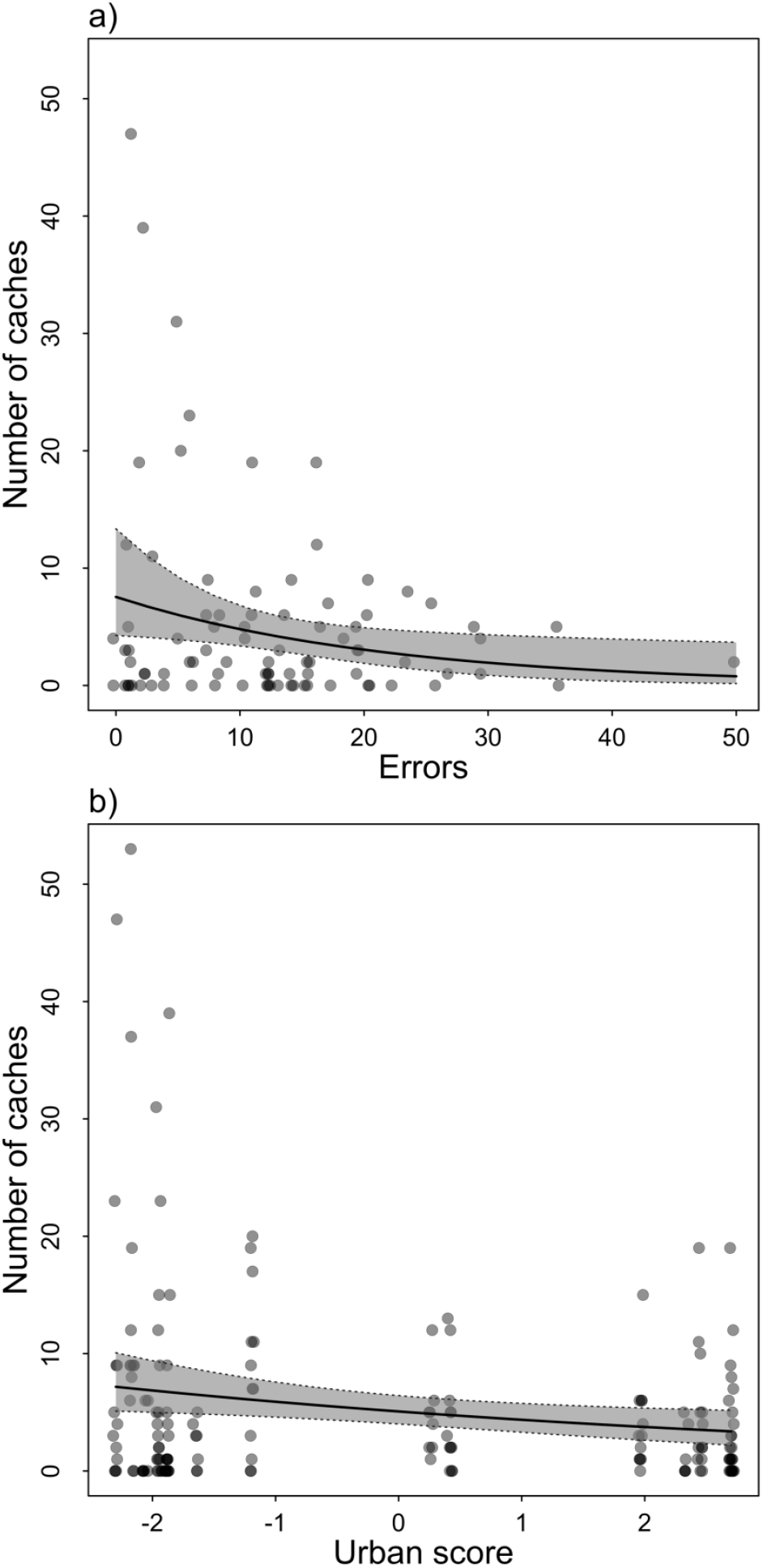
The relationship between the number of cage caches made by individuals in captivity and the a) number of errors made during the one-trial associative spatial task (*N* = 76), and b) urbanization score, where higher values represent more urbanized sites (*N* = 136). Significant effects and 95% confidence intervals from a generalized linear mixed model with a negative binomial distribution are shown. Points have been made partially transparent to show overlap.

**Table 1.** Predictors of the number of caches made by individuals in home cages fitted with a generalized linear-mixed effect model with random intercepts for sites in the (a) full hypothesized model, (b) full model once dropping the interaction, and (c) final model.

| Predictors | Estimate ±SE | $\chi^2$ | *P* |
| --- | --- | --- | --- |
| (a) Full model (N = 76) |  |  |  |
| Urban score | -0.33 ±0.43 | 0.81 | 0.37 |
| Errors on spatial task | -2.18 ±0.91 | 5.87 | 0.015 |
| Baseline corticosterone | 1.47 ±0.87 | 2.90 | 0.089 |
| Urban*Errors | 1.74 ±2.38 | 0.53 | 0.46 |
| (b) Dropped interaction (N = 76) |  |  |  |
| Urban score | -0.38 ±0.43 | 0.78 | 0.38 |
| Errors on spatial task | -2.20 ±0.92 | 5.71 | 0.017 |
| Baseline corticosterone | 2.07 ±1.27 | 2.65 | 0.10 |
| (c) Excluding spatial task data (N = 136) |  |  |  |
| Urban score | -0.75 ±0.31 | 6.06 | 0.014 |
| Baseline corticosterone | 1.14 ±0.77 | 2.60 | 0.14 |

### One-trial associative spatial memory task

Birds found the seed reward in the experimental phase of the spatial task with significantly fewer errors than would be expected by random searching in both sessions (chance = 30.5 following negative hypergeometric distribution; Tillé, Newman, & Healy, 1996) (Wilcoxon test: session 1, *N* = 96, median = 13, *P* < 0.001; session 2, *N* = 47, median = 21, *P* < 0.001). Urbanization and exploration scores were not found to be significant predictors of the number of errors or the latency to contact the seed reward in the experimental phase of the one-trial spatial task (Table 2c; Figure 2). Results were qualitatively unchanged when running spatial memory accuracy and latency models using only adult birds or when urbanization was assessed on alternative scales (200 m scale or as binary predictor; Table S2). Urbanization was still found to be a non-significant effect when directly comparing the number of errors between urban and rural habitats (Welch’s t-test: t = 1.38, df = 91.56, *P* = 0.17).

**Fig.2.**
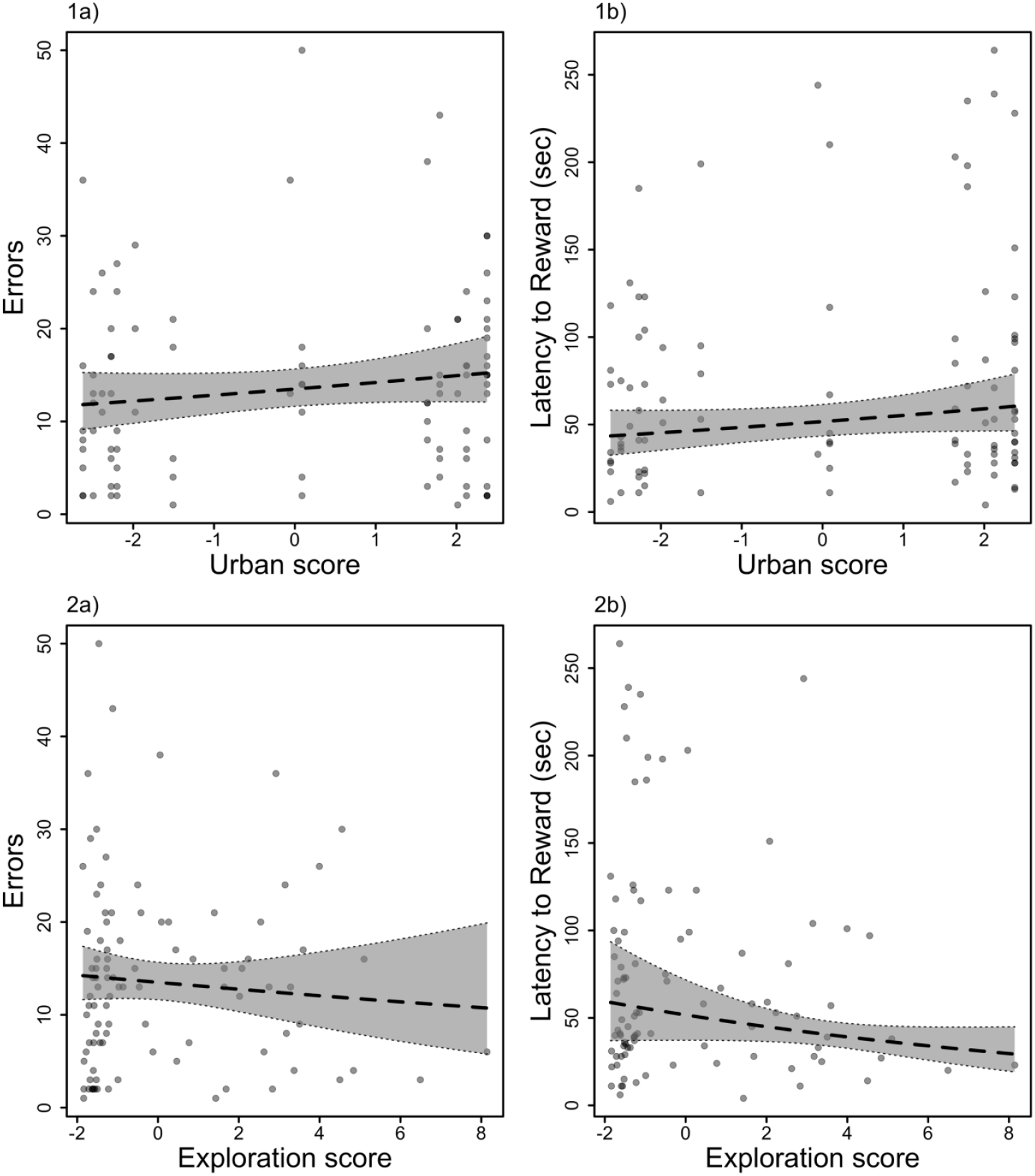
Final models showing the relationship between the effects of 1) urbanization and 2) individual exploration score, on the a) number of errors and b) latency to contact the hidden seed reward in the experimental phase of the one-trial associative spatial memory task. Non-significant estimated effects and corresponding 95% confidence intervals are shown. Points have been made partially transparent to show overlap.

**Table 2.** Predictors of the (1) number of errors (spatial memory accuracy) fitted using a generalized linear mixed-effects model with random intercepts for sites, and (2) latency to contact the reward fitted with a linear mixed-effects model with random intercepts for sites, in the one-trial spatial memory task. Shows tested predictors in the (a) full model, (b) full model once dropping the interaction, and (c) final model using the full dataset.

| Predictors | (1) Number of errors |  |  | (2) Latency to Reward |  |  |  |
| --- | --- | --- | --- | --- | --- | --- | --- |
| | Estimate ±SE | $\chi^2$ | *P* | Estimate ±SE | *F* | DF | *P* |
| (a) Full model (N=76) |  |  |  |  |  |  |  |
| Urban score | 0.41 ±0.21 | 3.64 | 0.056 | 0.41 ±0.25 | 2.50 | 1,5 | 0.18 |
| Exploration score | -0.49 ±0.46 | 1.09 | 0.30 | -0.77 ±0.52 | 2.63 | 1,69 | 0.11 |
| Order tested | 0.48 ±0.29 | 2.70 | 0.10 | 0.085 ±0.35 | 0.058 | 1,70 | 0.81 |
| Baseline corticosterone | -0.077 ±0.41 | 0.036 | 0.85 | -0.020 ±0.46 | 0.0020 | 1,70 | 0.96 |
| Urban*Exploration | 0.23 ±1.02 | 0.050 | 0.82 | -1.00 ±1.18 | 0.67 | 1,70 | 0.42 |
| (b) Dropped interaction (N=76) |  |  |  |  |  |  |  |
| Urban score | 0.40 ±0.21 | 3.62 | 0.057 | 0.42 ±0.25 | 2.37 | 1,5 | 0.19 |
| Exploration score | -0.46 ±0.44 | 1.08 | 0.30 | -0.85 ±0.51 | 2.61 | 1,70 | 0.11 |
| Order tested | 0.48 ±0.29 | 2.70 | 0.10 | 0.089 ±0.35 | 0.063 | 1,70 | 0.80 |
| Baseline corticosterone | -0.072 ±0.41 | 0.031 | 0.86 | -0.057 ±0.45 | 0.015 | 1,71 | 0.90 |
| (c) Final model (N=96) |  |  |  |  |  |  |  |
| Urban score | 0.25 ±0.20 | 1.66 | 0.20 | 0.33 ±0.22 | 1.93 | 1,7 | 0.21 |
| Exploration score | -0.28 ±0.37 | 0.56 | 0.45 | -0.70 ±0.42 | 2.72 | 1,90 | 0.10 |
| Order tested | 0.29 ±0.26 | 1.23 | 0.27 | -0.11 ±0.30 | 0.13 | 1,92 | 0.72 |

When including data from 2015 and from both sessions of the one-trial spatial memory task in 2016 (*N* = 96 individuals, 47 both sessions), urbanization and exploration scores still had no significant effect on the number of errors made (GLMM: urban estimate ±SE = 0.20 ± 0.17, χ^2^ = 1.34, *P* = 0.25; exploration estimate ±SE = -0.18 ± 0.30, χ^2^ = 0.34, *P* = 0.56). Birds that underwent both sessions in 2016 made significantly more errors in the second session (GLMM: session estimate ±SE = 0.45 ±0.14, χ^2^ = 10.39, *P* = 0.0013), and were not found to be consistent in the number of errors they made during the one-trial spatial memory task (R < 0.001, χ^2^ = 0, *P* = 1). Urbanization and exploration scores were also a non-significant predictors of the latency for individuals to contact the reward during both sessions (LMM: urban estimate ±SE = 0.23 ± 0.19, F_1,89_ = 1 43, *P* = 0.23; exploration estimate ±SE = -0.56 ± 0.35, F_1,82_ = 2.46, *P* = 0.12). The latency to contact the reward did not differ significantly between sessions (LMM: session estimate ±SE = 0.22 ± 0.17, F_1,86_ = 1.71, *P* = 0.19), and individuals were not repeatable in their latency to contact the reward between sessions (R < 0.001, χ^2^ = 0, *P* = 1).

## DISCUSSION

The adaptive specialization hypothesis (ASH) has been supported in scatter hoarders along elevational and latitudinal gradients (Pravosudov and Clayton 2002; Freas et al. 2012; Roth et al. 2012), and has only recently been examined along a gradient differing in the degree of urbanization (Kozlovsky et al. 2017). As well, the relationship between exploratory personality and spatial cognition has received little empirical attention and has never been explored within the same individual scatter hoarders. We find weak support that urban chickadees cache less than some rural chickadees, but report no differences in spatial memory accuracy between individuals along this gradient. These results suggest that the caching behaviours of some individuals, but not spatial memory, are affected by urbanization in our subpopulations, and we thus do not provide full support for ASH along an urban gradient. As well, we report no significant association between exploratory personality and spatial cognition at the individual level.

Caching intensity was significantly affected by urbanization as hypothesized, where rural chickadees cached more while in captivity in comparison to urban chickadees. However, we found that this relationship was driven by a few rural chickadees caching a lot more than urban chickadees, and excluding these individuals caused the effect to become non-significant. Since this effect is sensitive to extreme values, we are unable to conclude that rural scatter-hoarders cache more frequently than urban ones. Our results support findings from Kozlovsky et al. (2017) who found no differences in caching rates between city and forest mountain chickadees when measuring the number of caches made in a larger testing room. Individuals under captive conditions generally have access to *ad libitum* food sources, and it is therefore possible that longterm access to stable food sources may lead to downward adjustments in food hoarding. However, the current study measured caching intensity only a couple days after removal from the wild. Using this different approach, we thus provide support for findings from Kozlovsky et al. (2017) where mountain chickadees were held for longer-term captive conditions before testing. Rural chickadees may not cache more frequently than urban conspecifics in general, but may show more individual variation in their caching behaviours. Less stable and readily available resources in rural environments, in comparison to urban areas, may increase competition between chickadees in the same flock and cause individuals to adopt different foraging strategies. Dominance rank has previously been suggested as an individual trait that may affect caching intensity in scatter hoarders (Brodin et al. 2001). It is therefore possible that more dominant individuals cache less if they are able to monopolize food sources, causing less dominant individuals in their flocks to rely more on caching behaviours, with this effect being more pronounced in rural habitats.

We predicted that urbanization would be a significant predictor of spatial accuracy in the one-trial spatial memory task, and expected urban birds to perform more accurately than rural ones. However, we found no significant effect of urbanization on spatial accuracy or latency measures in the one-trial task. Captive baseline corticosterone was not found to significantly affect spatial performance measures and did not significantly differ between participating and non-participating individuals from each habitat type. Furthermore, the levels we measured shortly after the test are in the observed range for free-ranging black-capped chickadees in the wild (Montreuil-Spencer 2017) and after long-term captivity (Pravosudov et al. 2004). We therefore do not believe that stress levels after exposure to captivity impacted our conclusions on the link between urbanization and spatial memory performance. Interestingly, we found that individuals who performed more accurately on the spatial task also cached more frequently. Therefore, although individual differences in spatial performance were not predicted by habitat as hypothesized, we do find evidence for ASH at the level of the individual since birds that invested more in caching exhibited more accurate spatial memory. This result was sensitive to extreme values and we therefore conclude that some, but not all individuals, showed this association. This is the first evidence to our knowledge for ASH in scatter hoarders that occupy the same habitat. This finding suggests that variation in specialization of spatial memory in scatter hoarders may occur on several hierarchical levels, from species to populations and to individuals, and calls for further empirical examination.

Degree of urbanization, or food stability, may not alter spatial abilities within scatter hoarders if (i) there is sufficient gene flow occurring between habitats, and/or (ii) urbanization is too recent to detect a response to the evolutionary-novel selective pressures. Work examining ASH in scatter hoarders initially compared distinct populations that were separated by large geographical distances and were likely genetically distinct (Pravosudov and Clayton 2002; Roth et al. 2012). However, further work along an elevational gradient showed differences in spatial abilities between subpopulations that were located only 10 km away (Freas et al. 2012), which is in the range of distances between our urban and rural sites. In mountain chickadees, individuals at high and low elevations were found to show behavioural differences that are thought to limit dispersal between sites (Kozlovsky et al. 2014), but evidence for limited gene flow is lacking as these sites did not show genetic differentiation (Branch et al. 2017). In our system, we do not have information regarding genetic population structure, and thus gene flow may be acting as a buffer against divergent selection on spatial abilities, especially if selection against spatial abilities in urban environments is weak. Another potential explanation for the lack of relationship between urbanization and spatial memory is that the change in selection in urban areas may be too novel for an evolutionary response to have taken place. We did detect other behavioural differences along this gradient in our black-capped chickadee populations: individuals from more urbanized habitats are faster explorers (Thompson et al., in press) and seem to rely more heavily on social information about novel food patches (Jones et al. 2017). However it is not known at the moment if these differences are due to plasticity and/or micro-evolutionary change (Miranda 2017).

It is possible that the spatial abilities of chickadees would only be affected by urbanization as a result of phenotypic plasticity. Scatter hoarders deprived of caching opportunities were shown to have hippocampal volumes similar to non-scatter hoarders, and may lose enhanced spatial memory due to lack of experience (Clayton and Krebs 1994). Therefore, it is possible that urban individuals may become less reliant on spatial memory compared to rural individuals only once they experience habitat-specific demands for caching in their environment. To examine this possibility, we re-ran our analyses including only adult birds who had experienced at least a year within their environment, but our conclusions remained the same. We report no differences in spatial memory abilities between individuals occupying areas that differ in the degree of urbanization, either as a result of divergent selective pressures or phenotypic plasticity. We thus come to similar conclusions as Kozlovsky et al. (2017) and do not provide full support for ASH in scatter hoarders along an urban gradient.

Urban habitats have been considered more spatially complex environments (Griffin et al. 2017). The environmental complexity hypothesis proposes that enhanced cognitive mechanisms have evolved to allow individuals to collect, retain, and process more diverse information in heterogeneous environments (Godfrey-Smith 2001). For example, storm petrels living in a forest habitat were found to have larger relative hippocampi than their conspecifics living in an open meadow. The authors explain that these findings may be due to higher environmental complexity in forest habitats and speculate that individuals within the forest evolved enhanced spatial cognition for navigation purposes (Abbott et al. 1999). The spatial abilities of urban scatter-hoarders may thus not be lower than those of rural conspecifics if urban birds need to map a more spatially-complex environment. However, this explanation for a lack of difference in spatial memory is unlikely since we did not observe clear differences in caching intensity along our gradient. Although supplementary food sources have been shown to increase over-winter survival in chickadees (Brittingham and Temple 1988; Desrochers et al. 1988), artificially-elevated feeding regimes have not been found to create foraging dependency in this species (Brittingham and Temple 1992), suggesting the caching and foraging behaviours of chickadees may remain similar even when supplementary food is available.

Exploratory personality is hypothesized to affect the quality of information collected by individuals (Verbeek et al. 1994; Sih and Del Giudice 2012). We thus predicted that slow explorers would perform more accurately, and fast explorers would complete the spatial task more quickly. However, we report no significant interaction between exploratory personality and degree of urbanization, and no main effect of exploration, on spatial performance measures. Our study does not provide support for a trade-off between speed of exploration and spatial memory accuracy within a scatter hoarding model. As of yet, only a few studies have examined the relationship between spatial cognition and exploration in the same individuals, and results are mixed (mallards *Anas platyrhynchos*, Bousquet et al., 2015; lizards *Eulamprus quoyii*, Carazo, Noble, Chandrasoma, & Whiting, 2014; mice *Micromys minutus*, Schuster et al., 2017). Spatial memory is an important adaptive cognitive process used by scatter hoarders for survival, so perhaps this cognitive trait evolves independently of other traits, such as personality. Exploring whether a trade-off between these traits exists in non-hoarding species would be useful to understand whether our findings are scatter-hoarder specific.

We conducted a second session of the one-trial task in an attempt to evaluate the repeatability of spatial memory performance within individuals (Griffin et al. 2015). When moving the location of the rewarded site in the second session of the one-trial task, individuals still performed significantly more accurately than expected by random searching, but made significantly more errors than in the first session and were not found to be consistent in their spatial accuracy. Individuals were also not consistent in their latency to contact the reward between sessions, but latency measures did not differ between sessions. In the future, having longer-term intervals between sessions may help reduce memory interference (but see Urhan and Brodin 2015), or repeatability could be demonstrated using two different tasks. Demonstrating a positive relationship between individual performance on different tasks would suggest tasks are measuring the same trait and would thus be important also for establishing convergent validity (Carter et al. 2013). It is to be noted that previous studies have not attempted to quantify repeatability of spatial memory within individual scatter hoarders, and we know of only one published study reporting repeatability estimates for spatial learning and recognition (mice *Micromys minutus;* Schuster et al. 2017). There is ample evidence that individuals demonstrate consistent differences in behavioural styles or personality (Bell et al. 2009), but little is known about how individuals differ in their cognitive styles (Sih and Del Giudice 2012; Morand-Ferron 2017). Establishing the consistency of individual cognitive performance across contexts and over time will be an important step in linking cognitive measures to other individual traits such as personality (Griffin et al. 2015).

In conclusion, ASH was not supported along an urban gradient since urban birds did not show inferior spatial memory accuracy in comparison to rural birds. The lack of support for ASH along this gradient may be due to one of the listed explanations above, or perhaps a combination of them. Our data suggest, for the first time, that ASH may occur within individuals occupying the same environment; further work is needed to determine what other individual or ecological factors predict individual differences in the reliance on spatial memory and caching behaviours. We report no significant relationship between spatial memory accuracy and exploratory personality at both the population and individual levels, and thus are unable to provide evidence that slow explorers collect higher quality spatial information either within or between habitat types. Nonetheless, identifying predictive relationships between foraging strategies, personality traits, and cognitive abilities is an obvious direction for future research, which will help create a clearer picture for understanding adaptive individual variation in rapidly-changing environments.

## Acknowledgements

We would like to thank Celia Bodnar, Nicolas Bernier, Teri Jones, Dariya Quenneville, and Isabel Rojas-Ferrer for help in the field capturing chickadees. For help in captivity, we thank Sofia Karabatsos and Kayla Humphreys. Thank you to Dr. Frances Bonier and Shannon Smith for conducting blood sample assays for corticosterone and sex. We are grateful to the City of Ottawa, the National Capital Commission, and Nature Conservancy of Canada for granting permission to access field sites.

## Ethical Approval

All procedures performed in studies involving animals were in accordance with the ethical standards of the institution or practice at which the studies were conducted, Animal Use Protocol 1758.

